# HOCl-producing Electrochemical Bandage is Active in Murine Polymicrobial Wound Infection

**DOI:** 10.1101/2024.03.19.585100

**Authors:** Derek Fleming, Ibrahim Bozyel, Christina A. Koscianski, Dilara Ozdemir, Melissa J. Karau, Luz Cuello, Md Monzurul Islam Anoy, Suzanne Gelston, Audrey N. Schuetz, Kerryl E. Greenwood-Quaintance, Jayawant N. Mandrekar, Haluk Beyenal, Robin Patel

## Abstract

Wound infections, exacerbated by the prevalence of antibiotic-resistant bacterial pathogens, necessitate innovative antimicrobial approaches. Polymicrobial infections, often involving *Pseudomonas aeruginosa* and methicillin-resistant *Staphylococcus aureus* (MRSA), present formidable challenges due to biofilm formation and antibiotic resistance. Hypochlorous acid (HOCl), a potent antimicrobial agent produced naturally by the immune system, holds promise as an alternative therapy. An electrochemical bandage (e-bandage) that generates HOCl *in situ* was evaluated for treatment of murine wound biofilm infections containing both MRSA and *P. aeruginosa* with “difficult-to-treat” resistance. Previously, the HOCl-producing e-bandage was shown to reduce wound biofilms containing *P. aeruginosa* alone. Compared to non-polarized e-bandage (no HOCl production) and Tegaderm only controls, the polarized e-bandages reduced bacterial loads in wounds infected with MRSA plus *P. aeruginosa* (MRSA: vs Tegaderm only – 1.4 log_10_ CFU/g, p = 0.0015, vs. non-polarized – 1.1 log_10_ CFU/g, p = 0.026. *P. aeruginosa*: vs Tegaderm only – 1.6 log_10_ CFU/g, p = 0.0015, vs non-polarized – 1.6 log_10_ CFU/g, p = 0.0032), and MRSA alone (vs Tegaderm only – 1.3 log_10_ CFU/g, p = 0.0048, vs. non-polarized – 1.1 log_10_ CFU/g, p = 0.0048), without compromising wound healing or causing tissue toxicity. Addition of systemic antibiotics did not enhance the antimicrobial efficacy of e-bandages, highlighting their potential as standalone therapies. This study provides additional evidence for the HOCl-producing e-bandage as a novel antimicrobial strategy for managing wound infections, including in the context of antibiotic resistance and polymicrobial infections.

## Introduction

The emergence of bacteria that are resistant to antibiotics demands the investigation of new antimicrobial strategies. This is particularly critical in the context of wound infections. Studies suggest that almost 90% of wound samples may carry microorganisms with resistance to at least one antibiotic, with about 30% exhibiting resistance to six or more antibiotics.^1^ Among these, *Pseudomonas aeruginosa* is a Gram-negative pathogen that is intrinsically resistant to multiple antibiotics and prone to acquiring resistance,^2^ and methicillin-resistant *Staphylococcus aureus* (MRSA), are frequently identified as wound infection culprits.^3,4^

The presence of biofilms, communities of microorganisms protected by a complex matrix of polysaccharides, proteins, DNA, and other substances called extracellular polymeric substance (EPS), further enhances resistance in pathogens like *P. aeruginosa* and MRSA. Biofilm-related infections can be challenging to treat with existing therapies, hindering wound healing and causing persistent inflammation.^5,6^ In the United States, around 7 million patients suffer from chronic wounds annually, with approximately 60% of these wounds associated with microbial biofilms.^7,8^ Given the recalcitrance of chronic wound infections, and the common involvement of multi drug-resistant *P. aeruginosa* and MRSA, it is essential to develop new antibiofilm strategies that do not contribute to further antibiotic resistance.

Hypochlorous acid (HOCl) is a reactive oxygen species (ROS), naturally produced by phagocytes, that has potent antimicrobial properties.^9,10^ In past studies it has been shown that HOCl is broadly effective at killing both bacterial and fungal pathogens.^11–13^ A barrier to clinical use has been the inability to continuously deliver microbicidal, non-toxic concentrations to the infection site. In past studies, we developed an electrochemical platform for the *in situ* generation of HOCl. This platform was active against both bacterial and fungal biofilms *in vitro*, and against *P. aeruginosa in vivo* wound infections.^11–15^ Here, it is shown that an HOCl-producing electrochemical bandage (e-bandage), controlled by a miniature ‘wearable’ potentiostat, is effective in treating murine wound biofilm infections containing *P. aeruginosa* and MRSA together (and MRSA alone). The effectiveness of an HOCl-generating e-bandage was assessed on infections in mouse wounds. The assessment involved measuring the decrease in live bacteria within the wound, examining the progress of wound healing through the reduction of wound size, scoring of purulence reduction, analyzing tissue histopathology, and measuring levels of blood biochemistry markers and inflammatory cytokines. Additionally, the concentration of HOCl in the wound was measured, and scanning electron microscopy was conducted on excised wound biofilms to evaluate the treatment’s impact on the biofilm matrix and the integrity and abundance of the bacterial cells. Lastly, HOCl producing e-bandage treatment was compared with systemic antibiotic treatment, and the ability of e-bandage treatment to potentiate concurrently administered systemic antibiotics evaluated.

## Methods and Materials

### Electrochemical bandage

The e-bandage and wearable potentiostat have been previously described.^15–17^ Briefly, the e-bandage comprises two carbon fabric electrodes (Panex 30 PW-06, Zoltek Companies Inc., St. Louis, MO) with surfaces measuring 1.77 cm^2^ each for the working and counter electrodes, along with a silver/silver chloride (Ag/AgCl) wire serving as a quasi-reference electrode (QRE). A wearable potentiostat, powered by a 3-volt coin cell battery, maintains the operational potential of the working electrode at +1.5 V_Ag/AgCl_. Carbon fabric electrodes are separated by two layers of cotton fabric, with an additional layer placed over the counter electrode to aid in moisture retention. These layers are secured using silicone adhesive. The QRE is positioned between the cotton fabric layers separating the carbon electrodes. Titanium wires (TEMCo, Amazon.com, catalog #RW0524) with nylon sew-on caps (Dritz, Spartanburg, SC, item#85) connect to opposite ends of the e-bandage and link to the potentiostat. Under physiological conditions, polarization of the bandage leads to the generation of HOCl through these reactions:

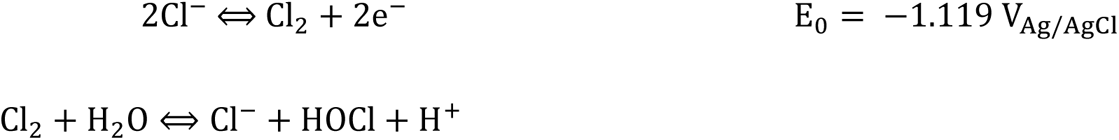

At pH 7.4 and 25°C, the conditions under which e-bandage was employed, HOCl dissociates to ∼57% HOCl and ∼43% ClO^-^.^18^

### Mice skin wound infection model

All animal experiments were approved by the Mayo Clinic Institutional Animal Care and Use Committee (A00003272-20). Full-thickness skin wounds were generated on Swiss Webster mice (Charles River, Wilmington, MA). Animals were anesthetized by intraperitoneal injection of a mixture of ketamine (90 mg/kg) and xylazine (10 mg/kg). Subcutaneous buprenorphine ER-Lab (1 mg/kg) was administered for analgesia. Creation of mature wound biofilms was as previously reported.^19,20^ The dorsal surface was shaved and disinfected, and a circular full-thickness skin wound created using a 5-mm biopsy punch (Acuderm Inc., Fort Lauderdale, FL). Wounds were then infected with 10 µl of 10^6^ colony-forming units (CFUs) of clinical isolates of MRSA IDRL-6169 and/or *P. aeruginosa* IDRL-11442 which has “difficult-to-treat” resistance, suspended in 0.9% sterile saline. MRSA IDRL-6169 is a methicillin and mupirocin-resistant isolate from a prosthetic hip. *P. aeruginosa* IDRL-11442 is a wound isolate resistant to piperacillin/tazobactam, cefepime, ceftazidime, meropenem, aztreonam, ciprofloxacin, and levofloxacin.^21^ Bacterial suspensions were permitted to settle in wound beds for 5 minutes. Subsequently, wounds were covered with semi-occlusive transparent Tegaderm® (3M, St. Paul, MN) secured using the liquid adhesive Mastisol® (Eloquest Health care, Ferndale, MI). Images of the wounds were captured, and wound diameters documented every other day using a Silhouette wound imaging system (Aranz Medical Ltd, Christchurch, NZ). Purulence was assessed before and after treatment to evaluate immune response to biofilm infection and treatment. The purulence scoring system was based on previous work^19^ using the following scale: 0 – no exudate in the wound-bed; 1 – slight turbid exudate at the wound site; 2 – mild amount of white exudate at the wound site; 3 – moderate amount of white exudate at the wound site; 4 – moderate amount of yellowish exudate at the wound site; 5 – large amount of turbid yellow exudate extending beyond the wound-bed.

### e-Bandage treatment

Following the establishment of 48-hours infections in mouse wound beds, mice were anesthetized with isoflurane, Tegaderm was removed, and wearable potentiostats were sutured to the scruff of the neck. Sterile e-bandages were pre-hydrated in sterile 1X phosphate-buffered saline (1X PBS), and 200 μL of sterile hydrogel (1.8% [w/v] xanthan gum in 1X PBS) were injected between the e-bandage layers. An additional 200 μL of hydrogel was applied to the wound beds, and e-bandages sutured on top to maintain close contact of the entire working electrode with the dorsal surface during mouse activity. e-bandages were then connected to the potentiostats and an additional 200 μL of hydrogel was placed on top, after which the entire e-bandage setups were covered with Tegaderm. Coin cell batteries (3V, Ecr1220 Energizer, St. Louis, MO) were inserted into the potentiostats to initiate e-bandage polarization (HOCl production). Treatment commerce for 48 hours with hydrogel refreshment and battery changes every 24 hours. Potentials of the working electrodes relative to the QREs were measured following treatment initiation, before and after each battery change, and prior to euthanasia to continuous operation.

Control groups included wounds administered only hydrogel and Tegaderm, and wounds treated with non-polarized e-bandages (i.e., no potentiostat or HOCl produciton). Additional animals from experimental and control groups underwent concurrent antibiotic dosing, with MRSA-infected mice treated with vancomycin and MRSA plus *P. aeruginosa*-infected mice treated with vancomycin and amikacin. Previously, the pharmacokinetic profiles of amikacin and vancomycin was established in Swiss Webster mice to determine a treatment dose of 15 mg/kg subcutaneous every 6 hours for amikacin and 150 mg/kg IP every 12 hours for vancomycin.^20^ At least 7 mice were included in each experimental and control group.

### Total wound HOCl measurement

Following wound bed excision and homogenization, the remaining portion (900 µL) of the wound homogenate, which was not used for quantifying bacterial load, was employed to determine the total wound HOCl content using free chlorine spectrophotometer test kits (TNT866; Hach Company, Ames, IA), following manufacturer’s instructions. In brief, homogenized wound contents were mixed with 4.1 mL of 1X PBS and centrifuged at 5000 rcf for 15 minutes. Resulting supernatants were filtered through syringe filters (0.22 µm pore size), and 4 mL of the filtrate added to free chlorine test tubes, allowing them to react for 1 minute before being measured at 515 nm using a Hach DR 1900 portable spectrophotometer (Hach Company). The free chlorine content was then converted to HOCl content using a specific equation, considering volume adjustment.

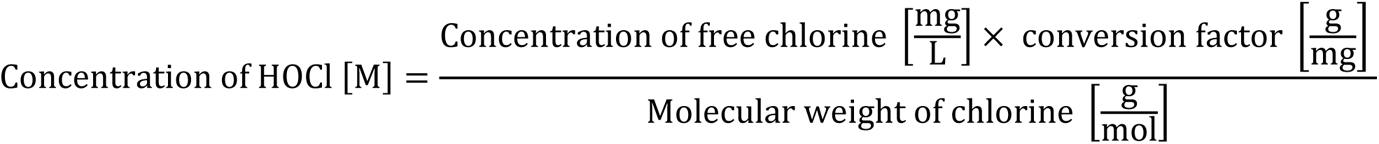

The molecular weight of chlorine (70.906 g/mol) and a conversion factor of 0.001 g/mg were used. Complete conversion of HOCl from free chlorine was assumed.

### Wound biofilm quantification

Following the conclusion of treatment, Tegaderm and e-bandages were removed from wound beds, and wound tissue excised using a 10 mm biopsy punch tool (Acuderm Inc., Fort Lauderdale, FL). Skin tissue was weighed, homogenized (Omni International, Kennesaw, GA) in sterile PBS, vortexed for 20 seconds and sonicated for 5 minutes in a water bath.

Subsequently, 100 µL of the resulting homogenate underwent serial dilution (10-fold dilutions) in 0.9% saline, and colony-forming units (CFUs) were determined by spread-plating 100 µL of each dilution onto tryptic soy agar with 5% sheep blood. Enumeration of mixed-species biofilm CFU counts was conducted using eosin methylene blue and colistin nalidixic acid agar plates. After 24 hours of incubation at 37°C, colonies were counted, and the results were reported as log_10_ CFU/g of tissue.

### Histopathology

For each treatment and control group, a subset of animals (n=3) was utilized for wound histopathology evaluation. The wounds were excised using a 10 mm biopsy punch and preserved in 10% formalin. After fixing, the specimens were dyed with hematoxylin and eosin (H&E) stains. Subsequently, a board-certified clinical pathologist, who was not aware of the sample origins, examined the slides. The pathologist assessed the level of inflammation on a scale from 0 (none) to 3 (severe), and checked for the presence of abscesses, ulceration, tissue death, and neutrophil infiltration, marking them as either present (Yes) or absent (No).

### Scanning Electron Microscopy

Following e-bandage treatment, the wound tissues from a subset of three animals from both the treatment and control groups were extracted with a 10 mm biopsy punch (Acuderm Inc., Fort Lauderdale, FL) and placed in sterile tubes containing a fixative solution composed of 4% formaldehyde plus 1% glutaraldehyde in phosphate buffer. The samples were then rinsed in PBS and dehydrated through a series of ethanol washes (10%, 30%, 50%, 70%, 90%, 95%, and 100% – twice). Dehydrated samples underwent critical point drying in a vacuum sputter coater (Bio-Rad E5100) and were coated with gold/palladium (60/40%). Finally, samples were visualized using a Hitachi S4700 cold-field emission scanning electron microscope (Hitachi High Technologies America, Inc., Schaumburg, IL). Samples assigned non-descriptive numbers upon collection by study staff and were then randomized by an electron microscopy technologist before imaging. Images were blindly reviewed by 8 members of the Mayo Clinic Infectious Diseases Research laboratory and scored on a scale of 1-3 for biofilm matrix integrity, bacterial cell integrity, and bacterial cell abundance.

### Toxicity screen analysis and inflammatory panel screening

After euthanasia, blood was drawn via cardiac puncture and then centrifuged to separate the serum. The serum samples were then examined for various biochemical markers using a Piccolo® Xpress™ Chemistry Analyzer at the Mayo Clinic Central Clinical Laboratory. This analysis included measuring levels of glucose, amylase, blood urea nitrogen, alkaline phosphatase, alanine aminotransferase, aspartate aminotransferase, gamma glutamyltransferase, lactate dehydrogenase, C-reactive protein, total bilirubin, creatinine, uric acid, albumin, total protein, calcium, chloride, magnesium, potassium, sodium, and total carbon dioxide. Furthermore, the serum was analyzed with a MesoScale Discovery SQ 120 to determine the presence of inflammatory biomarkers, including IFN-γ, IL-4, IL-5, IL-6, TNF-α, and KC/GRO.

### Statistical analysis

SEM scores from blind review were compared using ordinary two-way ANOVA with Tukey’s multiple comparisons test, with a single pooled variance. This allowed for comparison of the pooled reviewer scores for all sample types while accounting for reviewer and sample variability within each treatment group. For all other parameters, initial analysis among the experimental groups was performed using the Kruskal-Wallis test. For further detailed comparisons between specific groups, the Wilcoxon rank sum test was applied. The choice of non-parametric tests was driven by the small size of the samples and the lack of evidence supporting the normal distribution of the data. All statistical tests were conducted as two-tailed, considering p-values under 0.05 as statistically significant. When dealing with comparisons involving more than three groups, adjustments were made to account for the False Discovery Rate. The data analysis was conducted using SAS software (version 9.4, SAS Institute), while GraphPad Prism (version 10.1, GraphPad Software) was used for the creation of graphs.

## Results

### HOCl was produced by polarized e-bandages *in situ*

In previous studies, microelectrodes were used to demonstrate that e-bandages generate HOCl at the working electrode. HOCl was shown to penetrate biofilms, explant tissue, and wound beds in live mice.^15,16,22^ In this study, free chlorine spectrophotometer test kits were used to quantify the total concentration of HOCl in wounds infected with MRSA alone, and with *P. aeruginosa*. In both infection scenarios, wounds from mice treated with polarized electric bandages exhibited elevated levels of HOCl compared to those treated with non-polarized electric bandages or Tegaderm alone (**Figure 1**).

**Figure 1.**
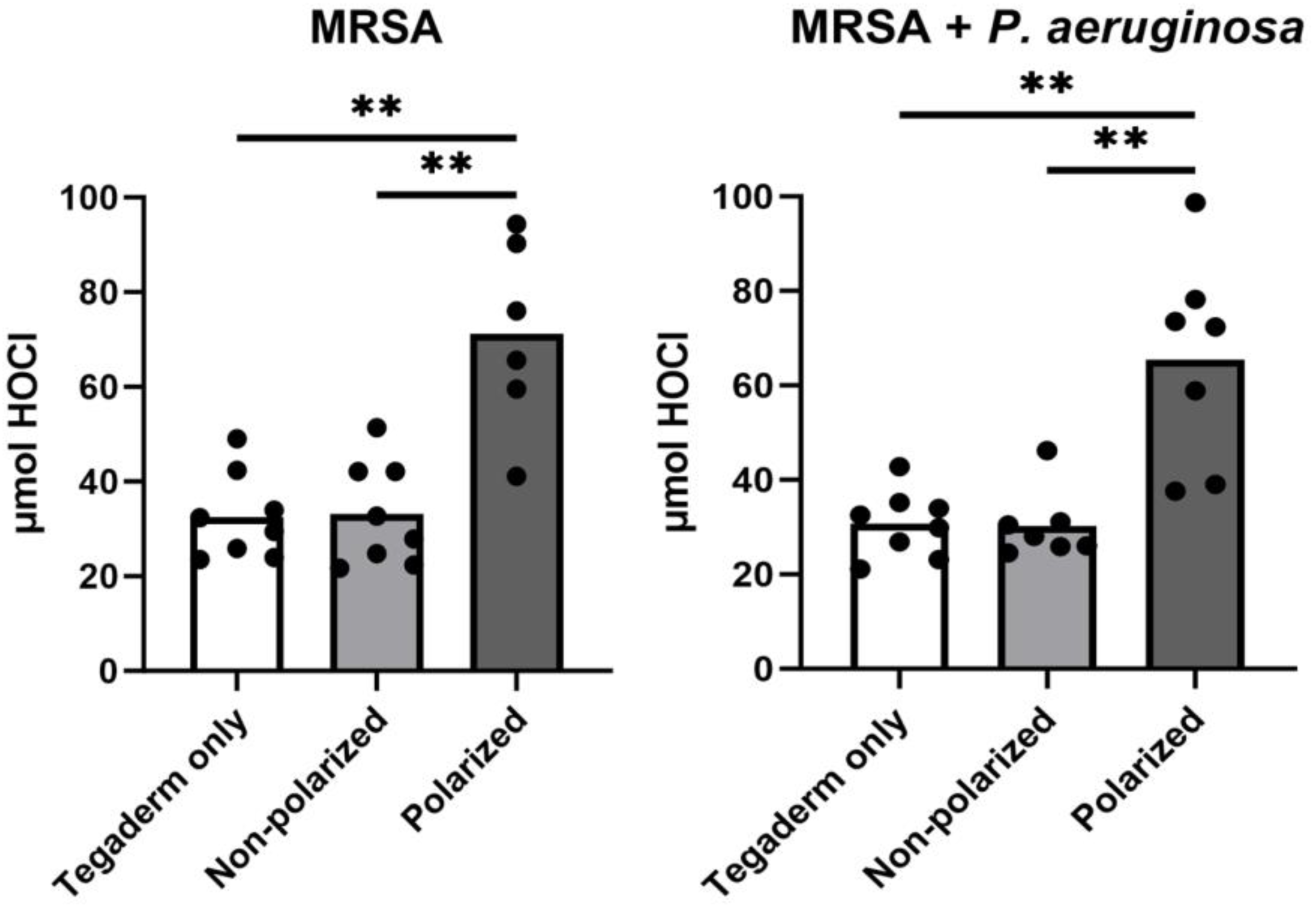
Polarized e-bandage treatment resulted in increased total wound HOCl content. 48-hour wound bed biofilms containing MRSA or MRSA plus *P. aeruginosa* (PA) were treated for 48 hours with either polarized (HOCl-producing) or non-polarized e-bandages and compared to Tegaderm only controls. Statistical analysis was performed using the Wilcoxon rank sum test with correction for false discovery rate. Individual data points with the means (bars) are shown. N ≥ 7. **p ≤ 0.01.

### Wound bacterial loads were reduced by polarized e-bandage treatment

To assess the efficacy of HOCl-generating e-bandage therapy in reducing bacterial biofilm burden *in vivo*, endpoint wound colony-forming units (CFUs) were quantified after 48 hours of treatment. Treatment of MRSA wound biofilms with polarized e-bandages reduced bacterial loads compared to non-polarized e-bandages (p = 0.0048) or Tegaderm alone (p = 0.0048, **Figure 2a**). Treatment of wound biofilms infected with both MRSA and *P. aeruginosa* by polarized e-bandages reduced bacterial loads of both species compared to non-polarized e-bandages (MRSA: p = 0.026; *P. aeruginosa*: p = 0.0032) or Tegaderm alone (MRSA: p = 0.003; *P. aeruginosa*: p = 0.0015; **Figure 2b**).

**Figure 2.**
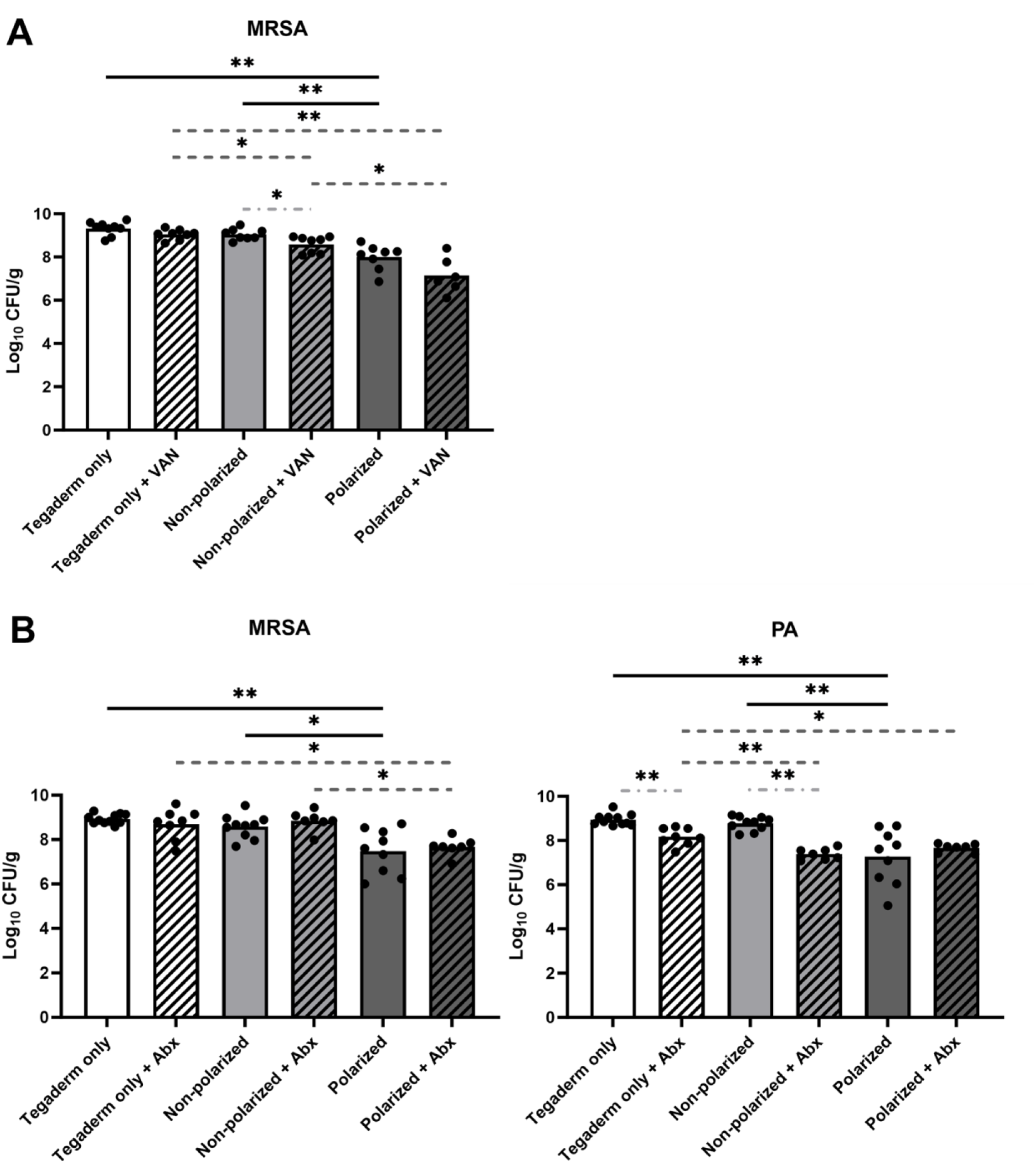
**Polarized e-bandage treatment reduces endpoint bacterial loads**. 48-hour wound bed biofilms containing MRSA (A) or MRSA plus *P. aeruginosa* (PA; B) were treated for 48 hours with either polarized (HOCl-producing) or non-polarized e-bandages, with or without systemic antibiotics (MRSA alone, vancomycin – VAN; MRSA plus *P. aeruginosa* – vancomycin plus amikacin – Abx) and compared to Tegaderm only controls, with and without antibiotics. Statistical analysis was performed using the Wilcoxon rank sum test with correction for false discovery rate. Individual data points with the means (bars) are shown. Solid black significance bars show differences between non-antibiotic-treated groups; Dashed dark grey significance bars show differences between antibiotic-treated groups; light grey dashed and dotted significance bars show differences between antibiotic and non-antibiotic-treated groups with the same e-bandage treatment type). N ≥ 7. *p ≤ 0.05, **p ≤ 0.01.

Three wounds from each treatment group for both MRSA alone and MRSA plus *P. aeruginosa* infected mice were blindly scored for biofilm matrix integrity, bacterial cell integrity, and bacterial cell abundance. Bacterial cell abundance was significantly lower after polarized e-bandage treatment for both MRSA and MRSA plus *P. aeruginosa*-infected wounds in comparison to Tegaderm only and non-polarized control groups (**Figure 3**), in agreement with reduced bacterial loads.

**Figure 3.**
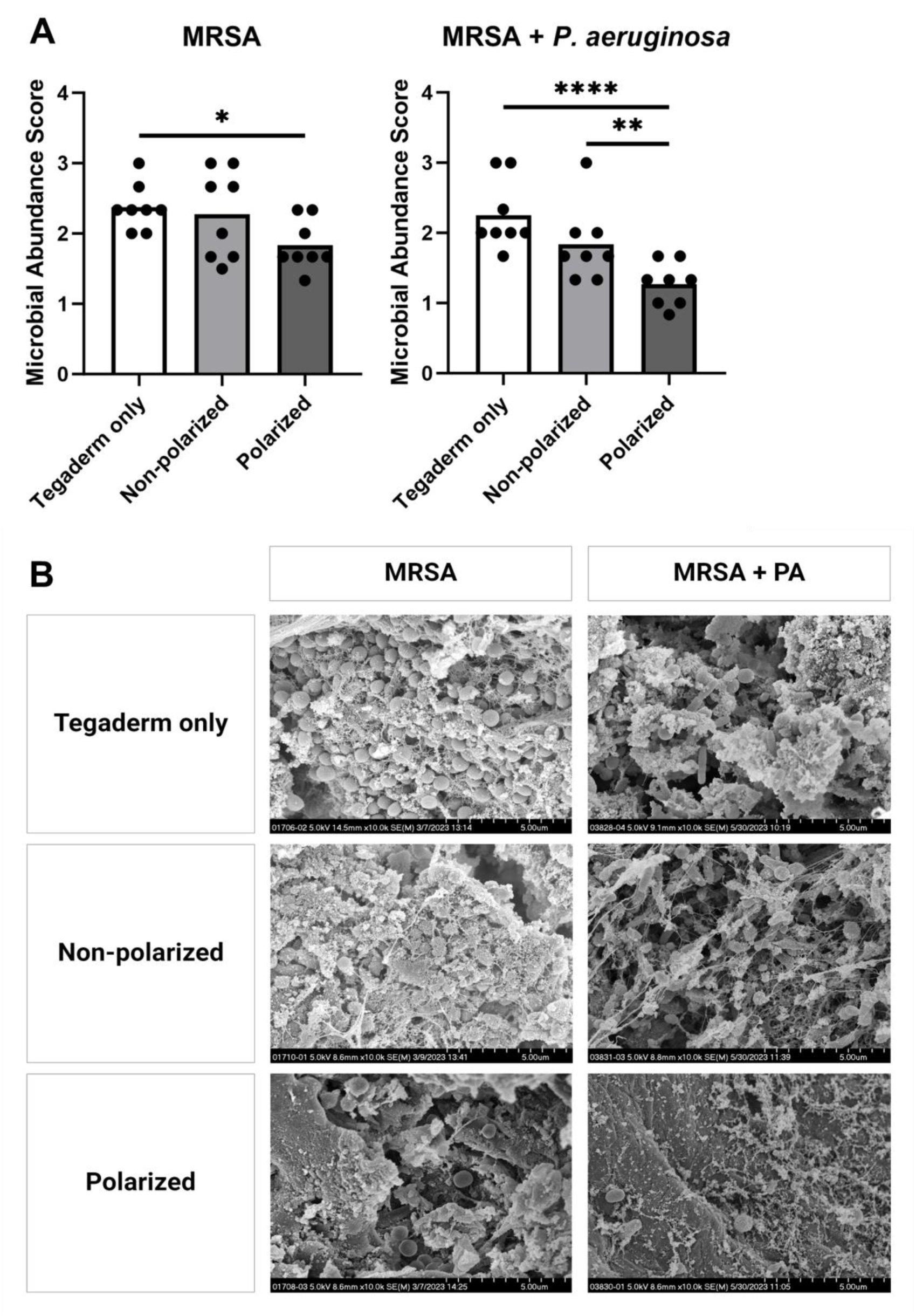
**Polarized e-bandage treatment reduces bacterial abundance observed in scanning electron microscopy (SEM) images**. SEM images of 48-hour wound biofilms containing MRSA or MRSA plus *P. aeruginosa* treated for 48 hours with polarized or non-polarized e-bandages, or Tegaderm only, were blindly reviewed and scored for bacterial abundance (A). Individual data points with the means (bars) are shown. Representative images are shown in (B) at 10,000 x magnification. Statistical significance was determined via two-way ANOVA with Tukey’s multiple comparisons test, with a single pooled variance. N = 3 samples per treatment type; 3-4 images per sample informed scoring. *p ≤ 0.05, **p ≤ 0.01. **p ≤ 0.0001.

To test if e-bandage treatment of established wound biofilms exhibited potential synergy with antibiotics against established wound biofilms, additional mice from all groups for both MRSA and MRSA plus *P. aeruginosa* infections were administered concurrent systemic vancomycin (for MRSA alone), or vancomycin and amikacin (for MRSA plus *P. aeruginosa*) for the duration of e-bandage treatment. Antibiotic treatment did not result in lower end-point bacterial loads for wounds treated with polarized e-bandages for either the single or dual-species infection groups. For MRSA alone, vancomycin reduced bacterial loads in only the non-polarized group (**Figure 2a**). For MRSA plus *P. aeruginosa*, MRSA load was not reduced in any group, however *P. aeruginosa* was reduced in both the Tegaderm only and non-polarized groups (**Figure 2b**).

### Wound healing was not hampered by polarized e-bandage treatment

To determine if HOCl-producing e-bandage treatment, with and without concurrent systemic antibiotics, affected wound closure over 48 hours of treatment, total wound area was measured before and after application. No significant differences in overall wound closure percentage were observed between any group for either MRSA alone, or MRSA plus *P. aeruginosa* infections (**Figure 4**). Interestingly, wound closure was less complete in the non-polarized group for MRSA plus *P.* aeruginosa-infected wounds when antibiotics were used.

**Figure 4.**
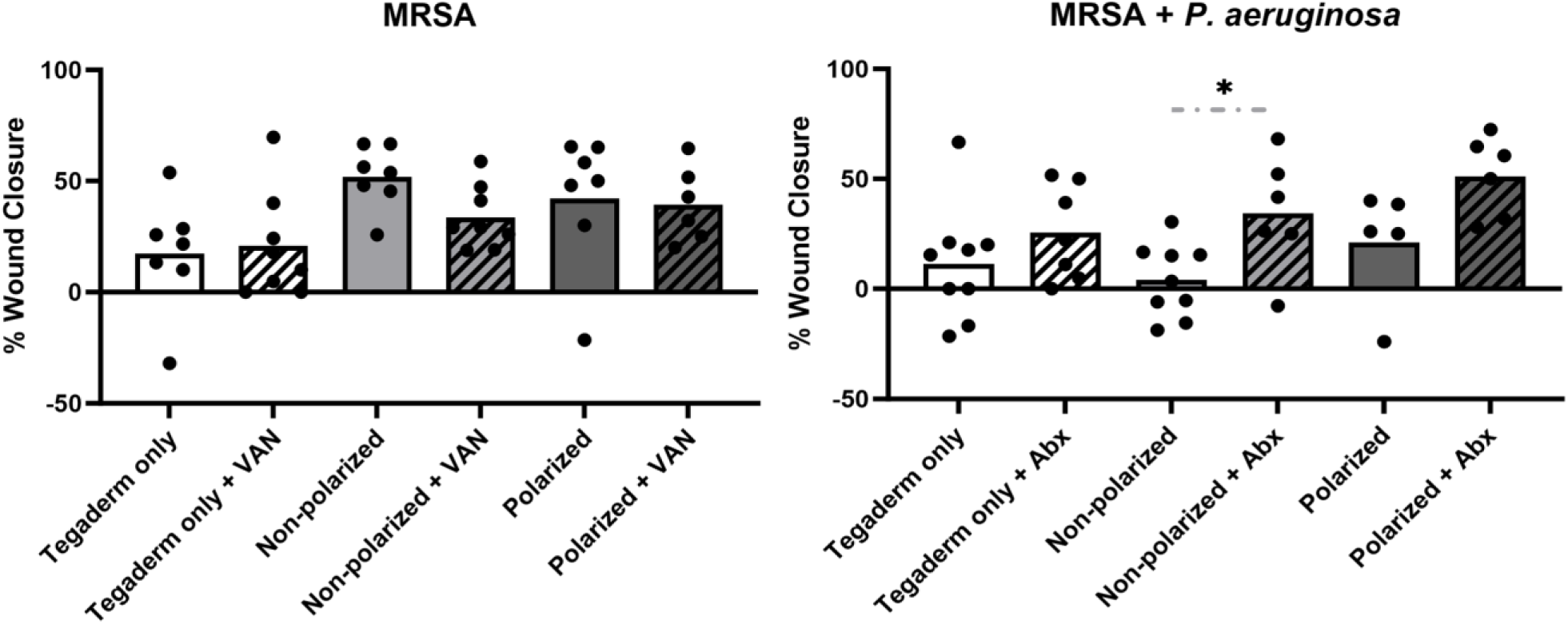
**Polarized e-bandage treatment did not hinder wound closure**. 48-hour wound bed biofilms containing MRSA (A) or MRSA plus *P. aeruginosa* (PA; B) were treated for 48 hours with either polarized (HOCl-producing) or non-polarized e-bandages, with or without systemic antibiotics (MRSA alone, vancomycin – VAN; MRSA plus *P. aeruginosa* – vancomycin plus amikacin – Abx) and compared to Tegaderm only controls, with and without antibiotics. Wound area was measured before and after treatment. Individual data points with means (bars) are shown. Statistical analysis was performed using the Wilcoxon rank sum test with correction for false discovery rate. N ≥ 7. *p ≤ 0.05.

This effect (though not significant) was also observed with the Tegaderm only and polarized groups for the dual-infection wounds, but with MRSA alone.

### Treatment of infected wounds with polarized e-bandages resulted in reduced purulence

The impact of e-bandage and/or antibiotic therapy on wound bed purulence was evaluated by scoring purulence before and after treatment (**Figure 5**). The use of polarized e-bandages resulted in a marked reduction in purulence compared to the Tegaderm-only control group in wounds infected with both MRSA and MRSA combined with *P. aeruginosa*. There was no significant improvement in purulence reduction between polarized and non-polarized e-bandage groups for either the mono or dual species infection, although the non-polarized group exhibited significantly less purulence than the polarized group in the dual-species infections (and to a lesser, insignificant amount in the MRSA only infections). Concurrent antibiotics did not improve purulence reduction in any treatment group in either the mono– or dual-species infected wounds.

**Figure 5.**
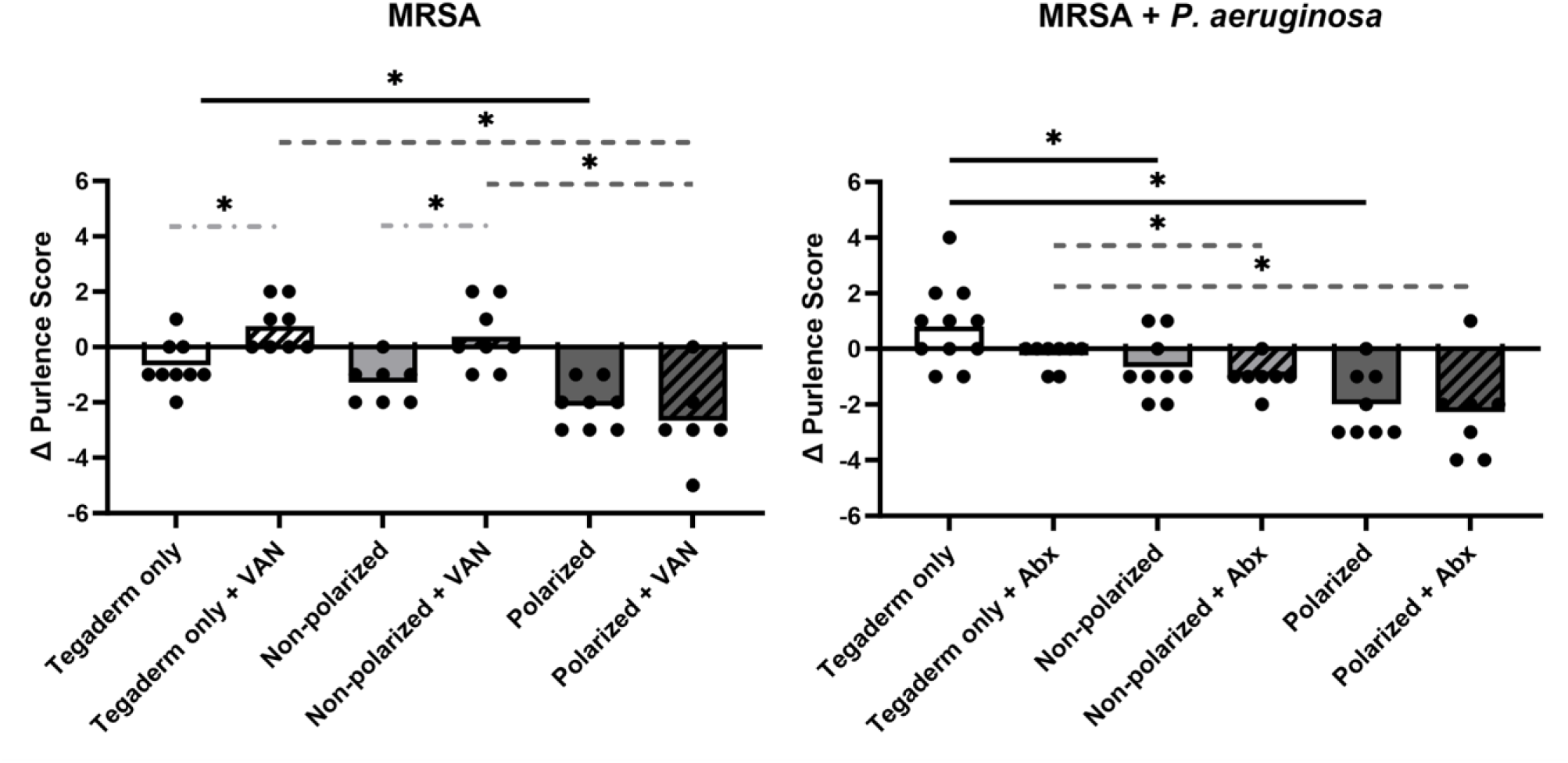
**e-Bandage treatment resulted in reduced wound purulence**. 48-hour wound bed biofilms containing MRSA or MRSA plus *P. aeruginosa* (PA) were treated for 48 hours with either polarized (HOCl-producing) or non-polarized e-bandages, with or without systemic antibiotics (MRSA alone, vancomycin – VAN; MRSA plus *P. aeruginosa* – vancomycin plus amikacin – Abx) and compared to Tegaderm only controls, with and without antibiotics. Wound purulence was scored before and after treatment. Statistical analysis was performed using the Wilcoxon rank sum test with correction for false discovery rate. Individual data points with means (bars) are shown. Solid black significance bars show differences between non-antibiotic-treated groups; Dashed dark grey significance bars show differences between antibiotic-treated groups; light grey dashed and dotted significance bars show differences between antibiotic and non-antibiotic-treated groups with the same e-bandage treatment type). N ≥ 7. *p ≤ 0.05.

### Polarized e-bandage treatment produced no observable tissue toxicity

To ascertain whether e-bandage therapy led to increased tissue toxicity compared to infection alone, samples were evaluated by a clinical pathologist blinded to the treatment. No notable variances were observed in overall inflammation, necrosis levels, abscess formation, ulceration, or neutrophilic inflammation across all treatment groups for both MRSA and MRSA plus *P. aeruginosa* infections.

### Assessment of inflammation and blood biomarkers for indication of animal health

Blood biochemical biomarker assessment and measurement of inflammatory cytokines was performed on a subset (n=3) of animals from each group to examine the immune response and general health of infected animals compared uninfected control animals at the time of euthanasia. As expected, all infected animals exhibited an elevated proinflammatory response compared to uninfected controls for both infection types (**Figure 6**). In particular, the proinflammatory cytokines INF-γ and IL-6 were elevated approximately 4 to16-fold and 2 to 9-fold respectively, indicating a strong, macrophage-driven immune response in all infected groups. Between infected groups, only KC/GRO showed significant elevation in animals treated with polarized vs non-polarized e-bandages. Notably, IL-6 was also most elevated in the polarized group for both infection types, albeit not to the level of statistical significance. For blood biochemical analysis, mean analyte levels were within normal healthy range for all groups, with no significant difference between animals treated with polarized or non-polarized e-bandages for both infection types (data not shown).

**Figure 6.**
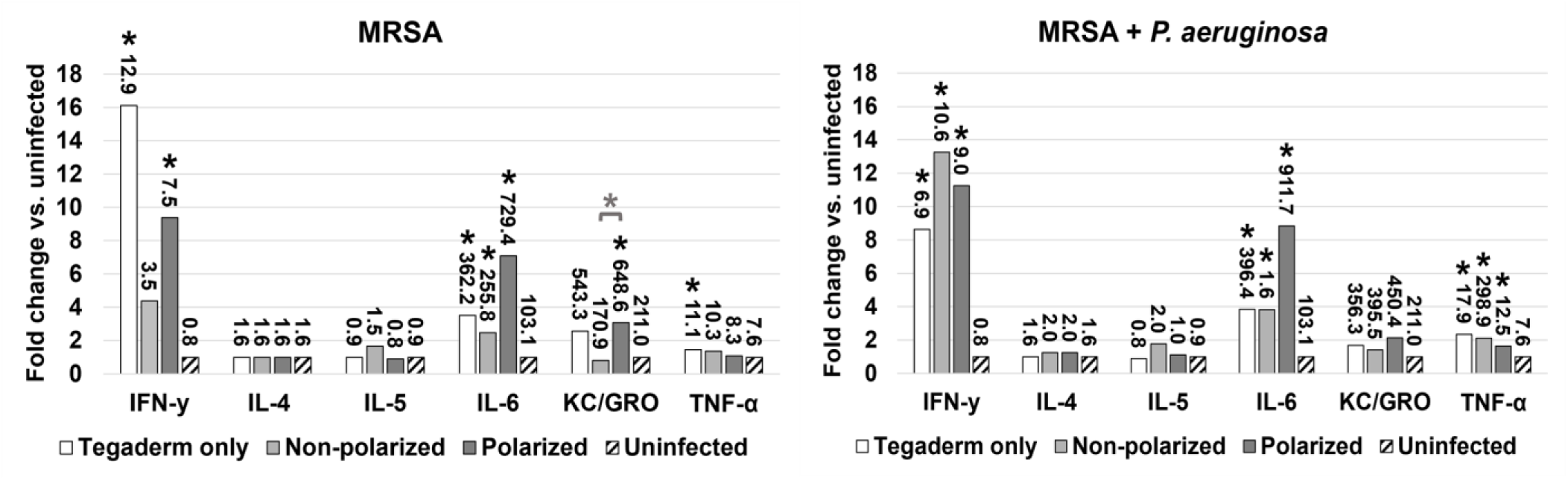
Inflammatory response in infected groups compared to uninfected controls. 48-hour wound bed biofilms containing MRSA or MRSA plus *P. aeruginosa* (PA) were treated for 48 hours with either polarized (HOCl-producing) or non-polarized e-bandages, with or without systemic antibiotics (MRSA alone, vancomycin – VAN; MRSA plus *P. aeruginosa* – vancomycin plus amikacin – Abx) and compared to Tegaderm only controls, with and without antibiotics. Following treatment, plasma collected and analyzed for levels of IFN-γ, IL-4, IL-5, IL-6, TNF-α, and KC/GRO. Fold change in comparison to uninfected controls is graphed, with analyte levels (in pg/ml) displayed as data labels. Statistical analysis was performed using the Wilcoxon rank sum test with correction for false discovery rate. Asterisks without bars represent significance compared to uninfected controls. Asterisks with bars represent significance between groups for the same analyte. N = 3. *p ≤0.05.

## Discussion

Development of alternative antimicrobial strategies is imperative in the face of rising antibiotic-resistant bacterial pathogens, particularly in the context of polymicrobial wound infections. In this study, efficacy of a previously developed HOCl-producing e-bandage for treatment of wound biofilm infections with antibiotic resistant clinical isolates of MRSA and *P. aeruginosa* was investigated. In a previous study efficacy of HOCl-producing e-bandages against wounds infected with *P. aeruginosa* alone was demonstrated.^15^ Polymicrobial infections, particularly with antibiotic resistant strains, pose additional challenges to wound infection healing. MRSA and *P. aeruginosa* are two of the most commonly isolated wound pathogens, and are often found together,^4,23^ with worse outcomes compared to mono-species infections.^24–26^

Results confirm the ability of polarized e-bandages to produce HOCl *in situ*, leading to elevated levels of HOCl in wound beds compared to non-polarized e-bandages or Tegaderm alone. Production of HOCl was associated with a significant reduction in bacterial biofilm burden *in vivo*, as demonstrated by lower bacterial loads in wounds infected with MRSA alone or co-infected with MRSA and *P. aeruginosa* following treatment with polarized e-bandages.

Further, blind review of SEM images of the wound beds taken from all groups revealed lower bacterial abundance in the animals treated with polarized e-bandages. No significant effect on the biofilm matrix was observed, indicating that the treatment is likely directly biocidal to biofilm-dwelling pathogens, as opposed to acting as an anti-EPS or pro-dispersal agent.

While polarized e-bandage treatment alone effectively reduced bacterial loads, addition of systemic antibiotics did not result in any additional microbicidal activity for either the MRSA infected or MRSA plus *P. aeruginosa* infected wounds, indicating that the antibacterial efficacy of e-bandages is independent of systemic antibiotic administration. This highlights the potential of e-bandages as a standalone antimicrobial strategy for wound infections, particularly in the context of antibiotic-resistant pathogens.

All infected groups showed an elevated, macrophage-driven immune response compared to uninfected controls. Between infected groups, animals treated with polarized e-bandages showed significantly elevated levels of KC/GRO when infected with MRSA alone, and insignificantly elevated levels of IL-6 when infected with both MRSA alone and in combination with *P. aeruginosa*. This indicates that inflammation in the polarized group may be more pronounced.

No adverse effects on wound healing or tissue toxicity associated with polarized e-bandage treatment was observed. Assessment of wound closure, purulence, histopathology, and blood biomarkers revealed no significant differences between non-polarized and polarized groups, indicating the safety and biocompatibility of e-bandage therapy in this context.

Previous results with e-bandages that produce an alternative reactive oxygen species, hydrogen peroxide (H_2_O_2_), found that wound healing was not only unimpeded but augmented,^20^ however, antimicrobial efficacy of electrochemically generated H_2_O_2_ was less than HOCl against a broad spectrum of microorganisms.^12,22,27–30^ Therefore, a programmable e-bandage that can produce both HOCl and H_2_O_2_ for optimal biocide and wound healing augmentation respectively should be explored.

In conclusion, these findings support promising efficacy of polarized HOCl-producing e-bandages in treating wound biofilm infections containing MRSA and *P. aeruginosa*. The ability of e-bandages to locally generate HOCl offers a novel and effective antimicrobial strategy that may address the challenges associated with antibiotic resistance in wound management, particularly in the context of polymicrobial infections. Further clinical studies are warranted to validate these findings and assess clinical application of e-bandage therapy for treatment of wound infections.

## Acknowledgments

Research reported in this publication was supported by the National Institute of Allergy and Infectious Diseases of the National Institutes of Health under award number R01AI091594. The content is solely the responsibility of the authors and does not necessarily represent the official views of the National Institutes of Health

## Disclosures

R.P. reports grants from MicuRx Pharmaceuticals and BIOFIRE. R.P. is a consultant to PhAST, Day Zero Diagnostics, Abbott Laboratories, Sysmex, DEEPULL DIAGNOSTICS, S.L., Netflix and CARB-X. In addition, R.P. has a patent on *Bordetella pertussis/parapertussis* PCR issued, a patent on a device/method for sonication with royalties paid by Samsung to Mayo Clinic, and a patent on an anti-biofilm substance issued. R.P. receives honoraria from Up-to-Date and the Infectious Diseases Board Review Course. H.B. holds a patent (US20180207301A1), “Electrochemical reduction or prevention of infections,” which refers to the electrochemical scaffold upon which the current design of e-bandage is based.

